# Relative contributions of the *ERG11*^VF125AL^ and *MRR1A*^N647T^ mutations to fluconazole resistance in Clade III *Candidozyma* (*Candida*) *auris* clinical isolates

**DOI:** 10.1101/2025.03.31.646430

**Authors:** Katherine S. Barker, Qing Zhang, Tracy L. Peters, Jeffrey M. Rybak, Joachim Morschhäuser, Christina A. Cuomo, P. David Rogers

**Author notes:** Corresponding Author: P. David Rogers, Pharm.D., Ph.D., Member and Chair St. Jude Endowed Chair in Pharmaceutical Sciences Department of Pharmacy and Pharmaceutical Sciences St. Jude Children’s Research Hospital 262 Danny Thomas Place Mail Stop 313, Memphis, TN 38105, t., f.

## Abstract

**Objectives:** *Candidozyma* (*Candida*) *auris* is an emerging fungal pathogen of global concern that often exhibits multi-drug resistance. Over 90% of isolates are resistant to fluconazole. Of the six described clades of *C. auris*, Clade III has been found to be nearly universally fluconazole resistant and almost every Clade III isolate described carries a mutation in the gene encoding the fluconazole target sterol demethylase (*ERG11*) leading to a VF125AL substitution and a mutation leading to a N647T substitution in the gene encoding Mrr1a, a transcriptional regulator of the Mdr1 transporter. Both mutations have been shown to contribute to fluconazole resistance in *C. auris*.

**Methods:** In the present study we introduced the Clade III *MRR1A* mutation into a Clade I background using CRISPR-Cas9 gene editing. In two Clade III clinical isolates we corrected the native *MRR1A* and *ERG11* mutations to their wild-type sequences as well as disrupted *MDR1*. Triazole susceptibilities and *MDR1* gene expression were measured in all strains.

**Results:** Introduction of the N647T substitution in a Clade I background confers a modest reduction in fluconazole and voriconazole susceptibility. Similarly, correction of *MRR1A* or disruption of *MDR1* in each Clade III background resulted in a one-dilution decrease in fluconazole and voriconazole MIC while the *ERG11* correction resulted in a three-dilution decrease in fluconazole and voriconazole MIC.

**Conclusions:** Our findings show that while the *MRR1A* mutation makes a modest contribution, the *ERG11* mutation is responsible for most of the fluconazole resistance observed in Clade III isolates. We also show that while these mutations likewise affect voriconazole susceptibility, they have no effect on susceptibility to itraconazole, isavuconazole, or posaconazole suggesting the potential therapeutic utility of these antifungals for infections due to Clade III isolates of *C. auris*.

## Introduction

*Candidozyma* (*Candida*) *auris* is a healthcare associated and multidrug-resistant pathogen of global concern associated with outbreaks of invasive candidiasis on multiple continents [1, 2]. It is the only fungal pathogen to be included in the urgent threat category of the U.S. Centers for Disease Control and Prevention (CDC) U.S. Antibiotic Threat Report [3]. Six clades of *C. auris* have been identified with Clades I, III, and IV causing the greatest number of infections at present. Over 90% of isolates are resistant to fluconazole, approximately 50% are resistant to amphotericin B, and approximately 5% are resistant to the echinocandins [4]. Fluconazole resistance has been shown to be due in part to mutations in the gene encoding the drug target, sterol demethylase (*ERG11*) or mutations in the gene encoding the transcriptional regulator *TAC1B* leading to overexpression of *CDR1* which encodes an ATP Binding Cassette (ABC) transporter homologous to similar transporters that contribute to resistance in other *Candida* species [5–8].

In *Candida albicans*, *MDR1* encodes a transporter of the Major Facilitator Superfamily that is upregulated in some fluconazole-resistant clinical isolates and contributes to this phenotype. Its overexpression is due to activating mutations in a transcription factor that we previously identified and named Mrr1 for <u>m</u>ultidrug resistance regulator. Introduction of activating mutations in *MRR1* into a susceptible strain confers increased fluconazole resistance and overexpression of *MDR1* [9]. The *C. auris* genome contains three genes encoding homologs of *C. albicans MRR1* [10]. Nearly all known isolates of *C. auris* Clade III are resistant to fluconazole and have been observed to carry a mutation in *MRR1A* leading to the N647T amino acid substitution [10, 11]. This mutation has been shown to contribute to fluconazole resistance as deletion in a Clade III isolate decreased the MIC from 512 µg/mL to 256 µg/mL and introduction of this mutation into a susceptible Clade IV isolate increased the MIC from 4 µg/mL to 16 µg/mL [12].

In addition to the *MRR1A* mutation leading to the A647T substitution (*MRR1A*^N647T^), almost all Clade III isolates also carry a mutation in *ERG11* leading to the VF125AL substitution (*ERG11*^VF125AL^) that has been shown to contribute to fluconazole resistance [6, 11]. This has complicated the assessment of the contribution of this *MRR1A* mutation to resistance in Clade III isolates. In the present study we further examined the effect of this *MRR1A* mutation and show that its contribution to fluconazole resistance in Clade III isolates is minimal with the majority of resistance being due to the mutation in *ERG11*. Moreover, we show that neither of these mutations have an effect on the susceptibility of Clade III isolates to isavuconazole, itraconazole, or posaconazole, highlighting the possible utility of these antifungals in the treatment of infections due to such isolates.

## Methods

### Cell growth conditions

*C. auris* strains and isolates were routinely propagated in YPD (1% yeast extract, 2% peptone, 2% dextrose) at 35°C and stored in 40% glycerol at –80°C.

### Strain construction

The clinical isolates and strains used in this study are listed in the web-only **Supplementary Table S1**, and the oligonucleotides used are listed in web-only **Supplementary Table S2**. The strains were constructed as described previously [8], with *MDR1* disruption and *ERG11* correction repair templates generated by PCR amplification of their corresponding gBlock oligonucleotides and the *MRR1A* correction repair template generated by primer extension.

### Minimum inhibitory concentration (MIC) determination

MICs for fluconazole, itraconazole, voriconazole, posaconazole, and isavuconazole were measured by modified CLSI broth microdilution assays as described previously [6]. Additionally, fluconazole Etests (Biomerieux) were performed as described previously [6].

### RNA isolation and qRT-PCR

An aliquot of cells from an overnight culture was used to inoculate 10 ml YPD to a OD_600_=0.08-0.12, followed by incubation at 35°C in a shaking incubator (220 rpm) until mid-log phase (6 hrs). Cell cultures were grown in triplicate. Cells were collected by centrifugation, supernatants removed, and cell pellets stored at –80°C. RNA was extracted using methods described previously with some modification [13]. Cell pellets were resuspended in 100 µl FE Buffer (98% formamide, 0.01 M EDTA) at room temperature. Fifty microliters of 1 mm RNase-free glass beads were added to the cells and were subjected to vortex disruption for 5 min at room temperature followed by snap-cooling on ice. The cell lysate was clarified by centrifugation, and the crude RNA extract applied to an RNeasy Qiagen column followed by washes and DNase treatment using the Qiagen RNeasy kit per manufacturer’s instructions. RNA integrity was confirmed by agarose gel electrophoresis, and concentrations were approximated by Nanodrop. cDNA was synthesized from 500 ng RNA using the RevertAid First Strand cDNA Synthesis Kit (Invitrogen/Thermo Fisher) with the provided random primer mix according to the manufacturer’s instructions. qRT-PCR was performed from three biological replicates, each with three technical replicates, using SYBR Green PCR master mix (Bio-Rad). Fold changes were calculated using the ΔΔ^CT^ method with *MDR1* CT values normalized to *ACT1* CT values (generating dCT values). Fold change calculations were performed by using the corresponding dCT value for each of the three biological replicates from 1c, 1102MRR1A-23A, and 384MRR1A-39E (for the 1c, AR1102, and AR0384 strains sets, respectively) as the comparator. Statistical significance was analyzed by one-way ANOVA.

## Results

### *MRR1A*^N647T^ confers a slight decrease in fluconazole susceptibility when expressed in a Clade I strain

To assess the impact of *MRR1A*^N647T^ on susceptibility, we first introduced it into a fluconazole-susceptible Clade I strain 1c which carries the wild-type *MRR1A* allele. Strain 1c is derived from highly fluconazole-resistant Clade I clinical isolate Kw2999 (fluconazole MIC = 256 µg/mL) that carries a mutation in *ERG11* leading to the K143R substitution and the mutation in *TAC1B* leading to the A640V substitution [14]. In strain 1c, both of these mutations have been corrected to their wild-type sequences using CRISPR-Cas9 gene editing resulting in a fluconazole MIC of 2 µg/mL [8].

Introduction of *MRR1A*^N647T^ into strain 1c resulted in approximately a 6-fold increase in *MDR1* expression (**Figure 1**) and a slight decrease in fluconazole susceptibility from 2 µg/mL to 4 µg/mL by both broth microdilution and Etest (**Figure 2**). A similar increase in susceptibility was observed for voriconazole but not for itraconazole, posaconazole, or isavuconazole (**Figure 3**).

**Figure 1.**
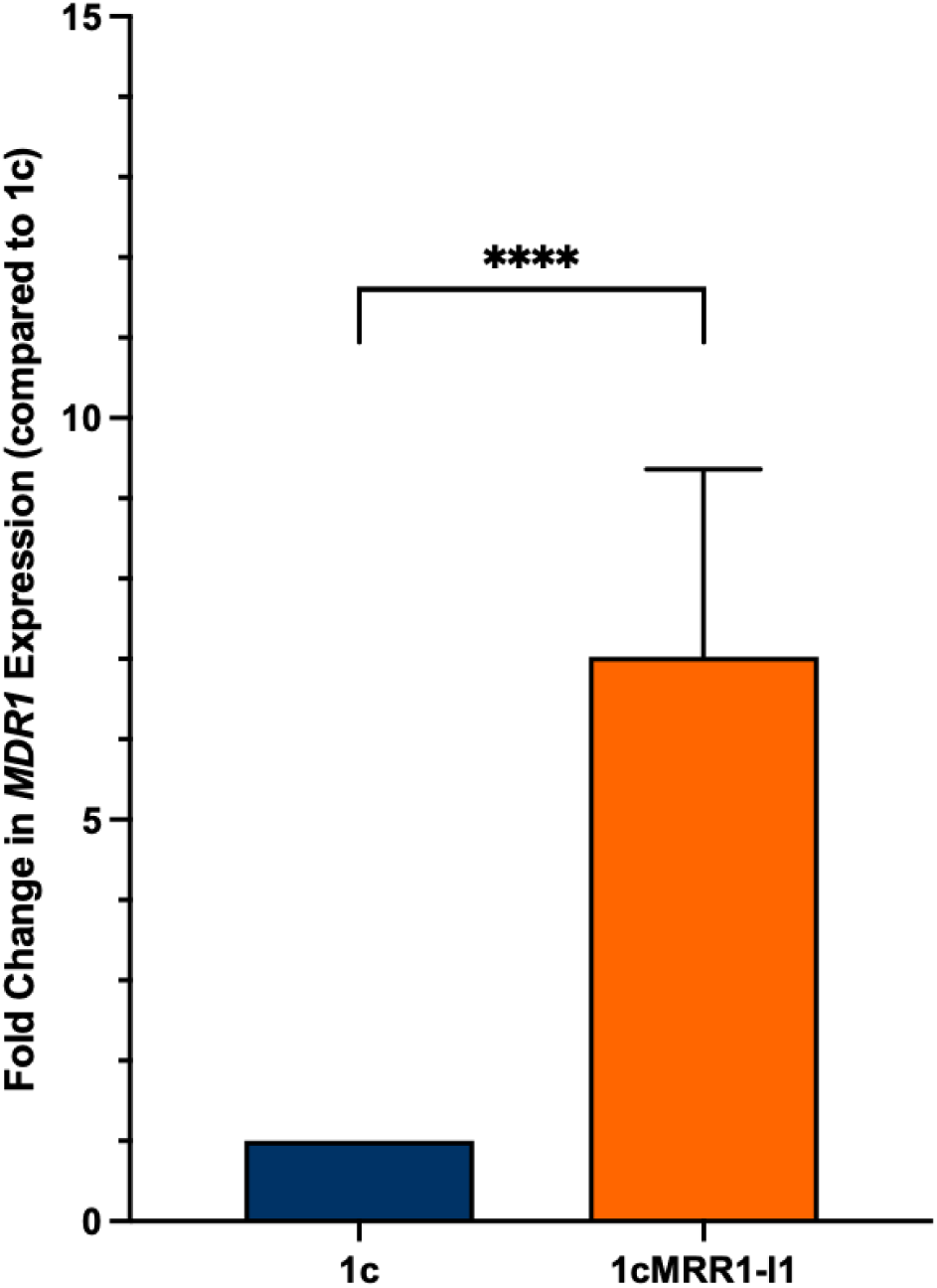
Fold change in *MDR1* gene expression in Clade I strain 1c (*MRR1A*^WT^) and strain 1cMRR1-I1 (*MRR1A*^N647T^). Fold changes were calculated as compared to the corresponding dCT value of 1c for each biological replicate. Asterisks (****) indicate *p*-value <0.0001.

**Figure 2.**
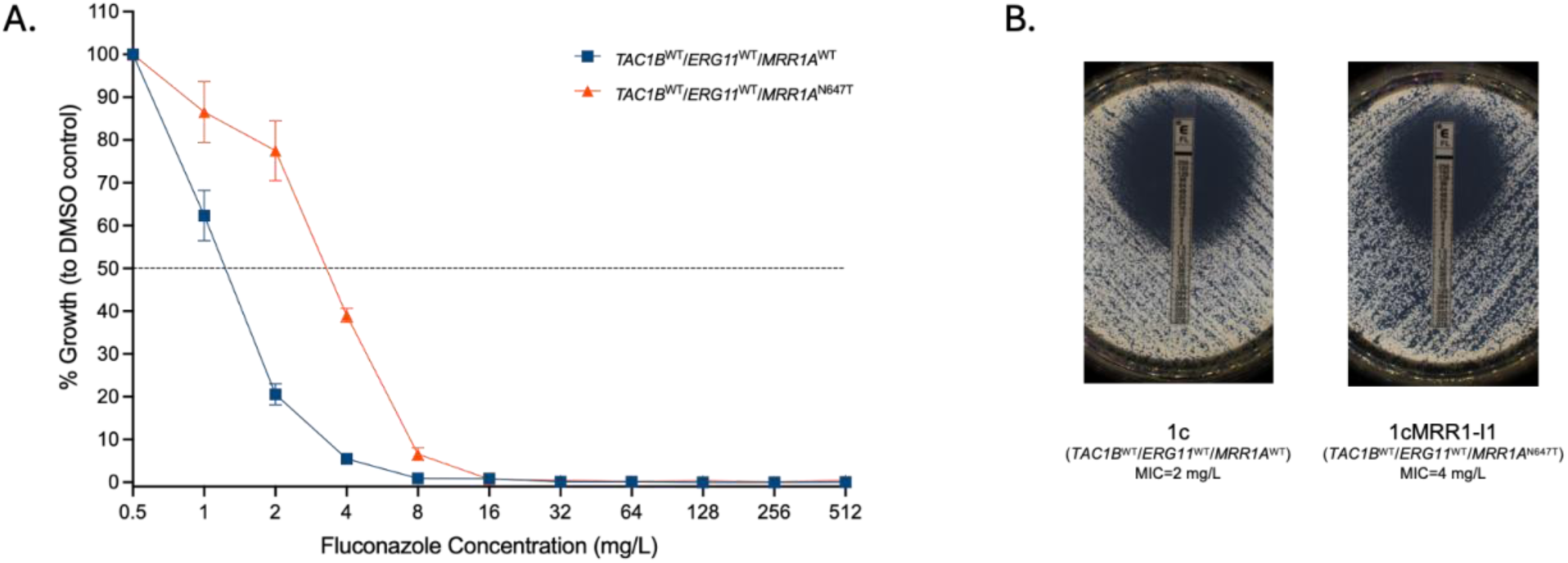
Fluconazole MIC of Clade I strain 1c and strain 1cMRR1-I1 (A) by broth microdilution and (B) by Etest.

**Figure 3.**
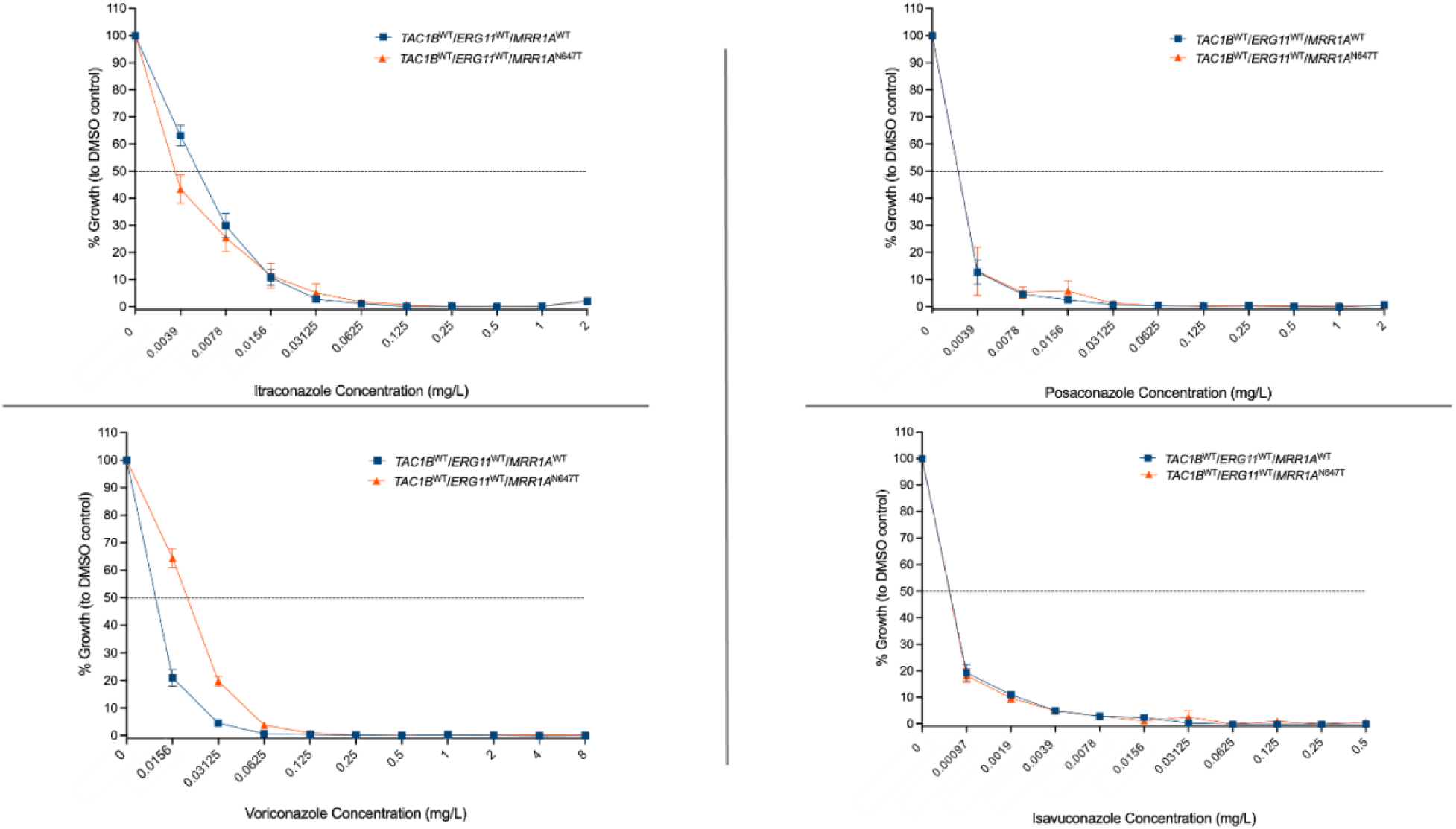
Itraconazole, posaconazole, voriconazole, and isavuconazole MICs of Clade I strains 1c and 1cMRR1-I1 as determined by broth microdilution.

### *ERG11*^VF125AL^ and to a lesser extent *MRR1A*^N647T^ drive fluconazole resistance in Clade III clinical isolates

In order to determine the contribution of *MRR1A*^N647T^ to fluconazole resistance in a Clade III background, we first edited this mutation to the wild-type sequence in two fluconazole-resistant Clade III clinical isolates, AR1102 and AR0384. This resulted in a 12– and nine-fold reduction in *MDR1* expression in these isolates (**Figure 4a and 4b**), respectively, and a reduction in fluconazole MIC from 64 µg/mL to 32 µg/mL by broth microdilution and no appreciable change by Etest in both isolates (**Figure 4c and 4d**). A similar increase in susceptibility was observed for voriconazole by broth microdilution, but not for itraconazole, posaconazole, and isavuconazole (**Figure 5a and 5b**).

**Figure 4.**
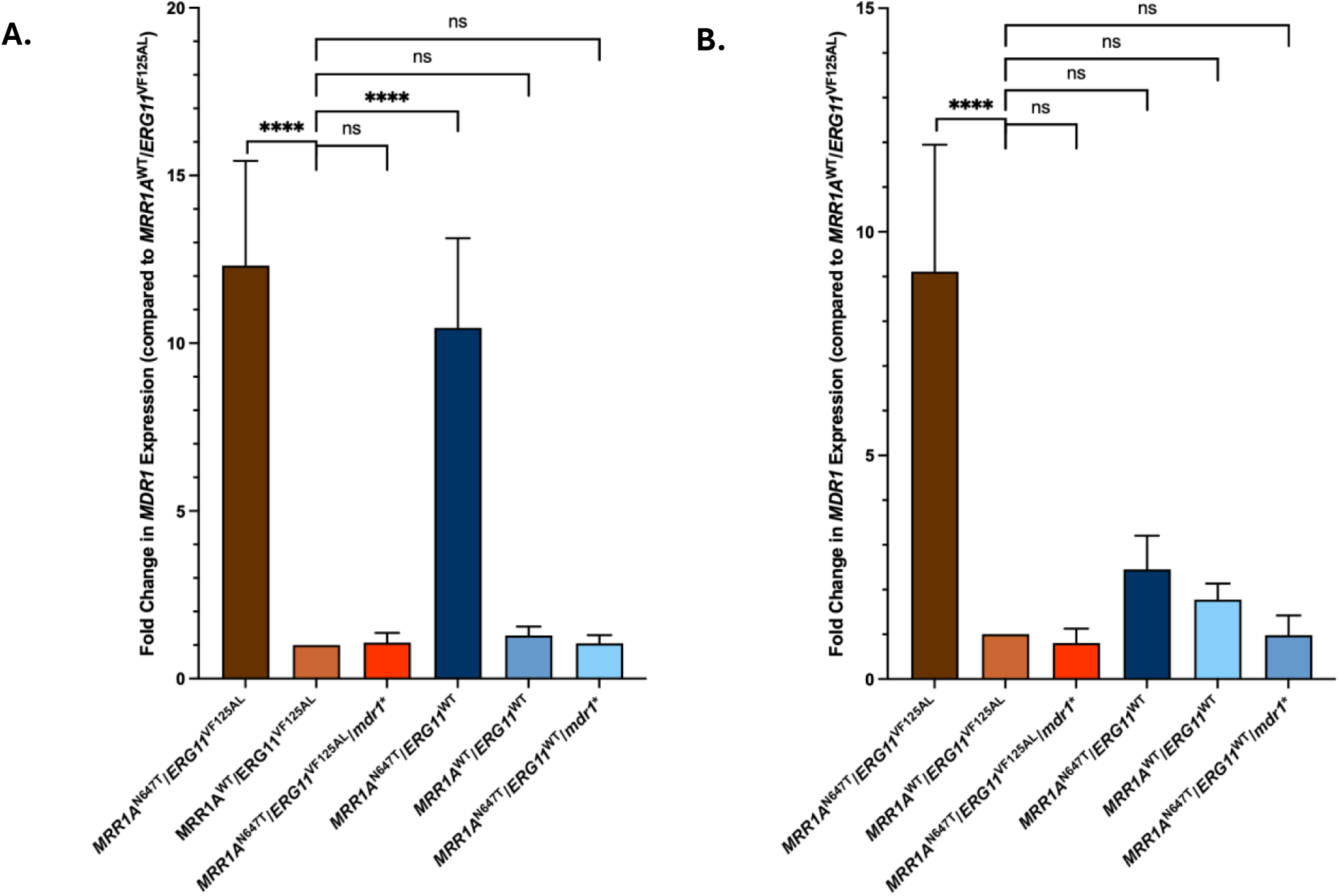
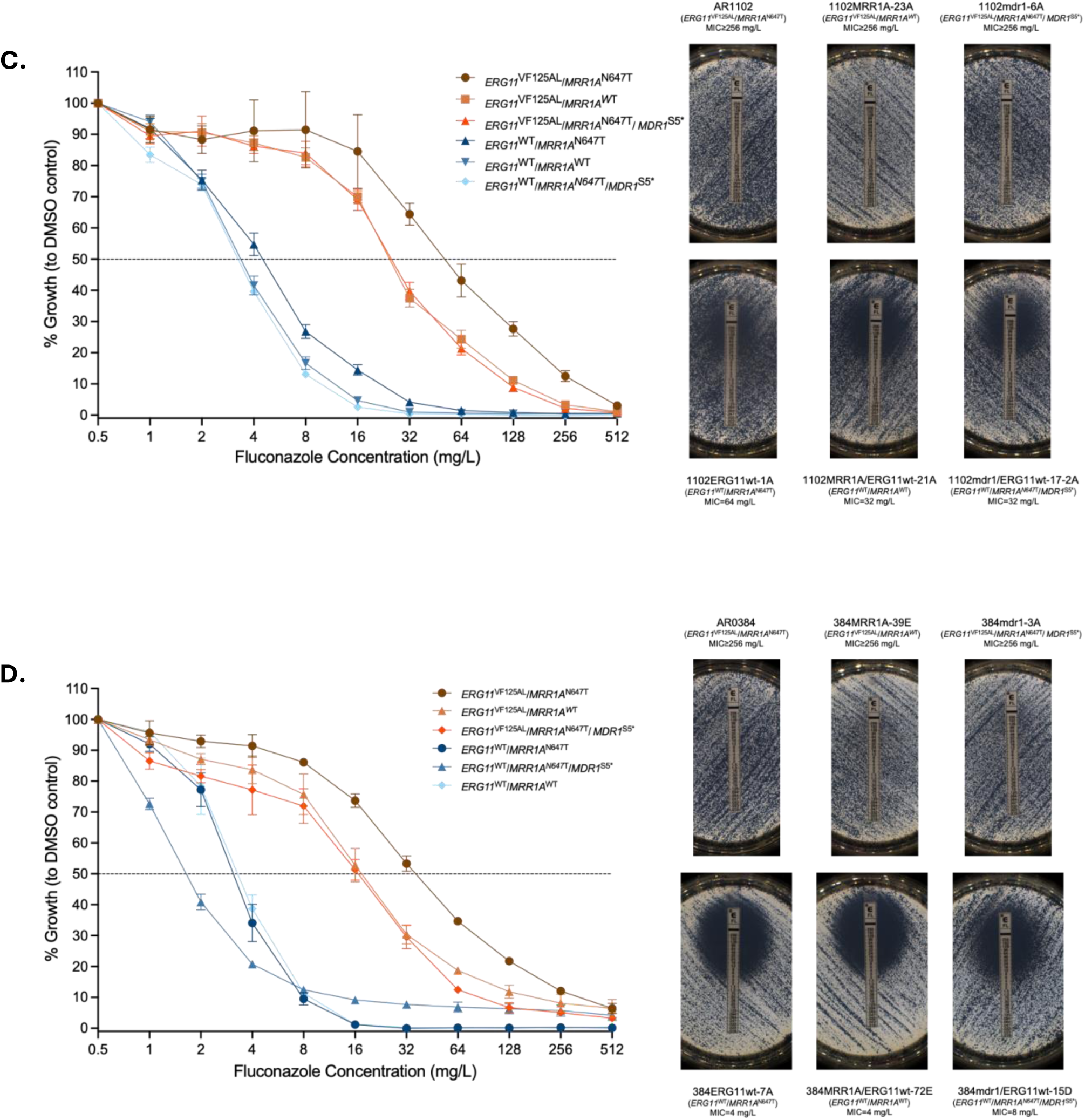
Fold change in *MDR1* gene expression in Clade III isolate AR1102 and its derivative strains (A) and Clade III isolate AR0384 and its derivative strains (B) and the corresponding FLC MIC determinations for the AR1102 strain set (C) and the AR0384 strain set (D). Fold changes in panel A were calculated as compared to the corresponding dCT value of 1102MRR1A-23A (*MRR1A*^WT^/*ERG11*^VF125AL^) for each biological replicate, and fold changes in panel B were calculated as compared to the corresponding dCT value of 384MRR1A-39E (*MRR1A*^WT^/*ERG11*^VF125AL^) for each biological replicate. Asterisks (****) indicate *p*-values of <0.0001. Panels C and D are comprised of both FLC broth microdilution and FLC Etests.

**Figure 5.**
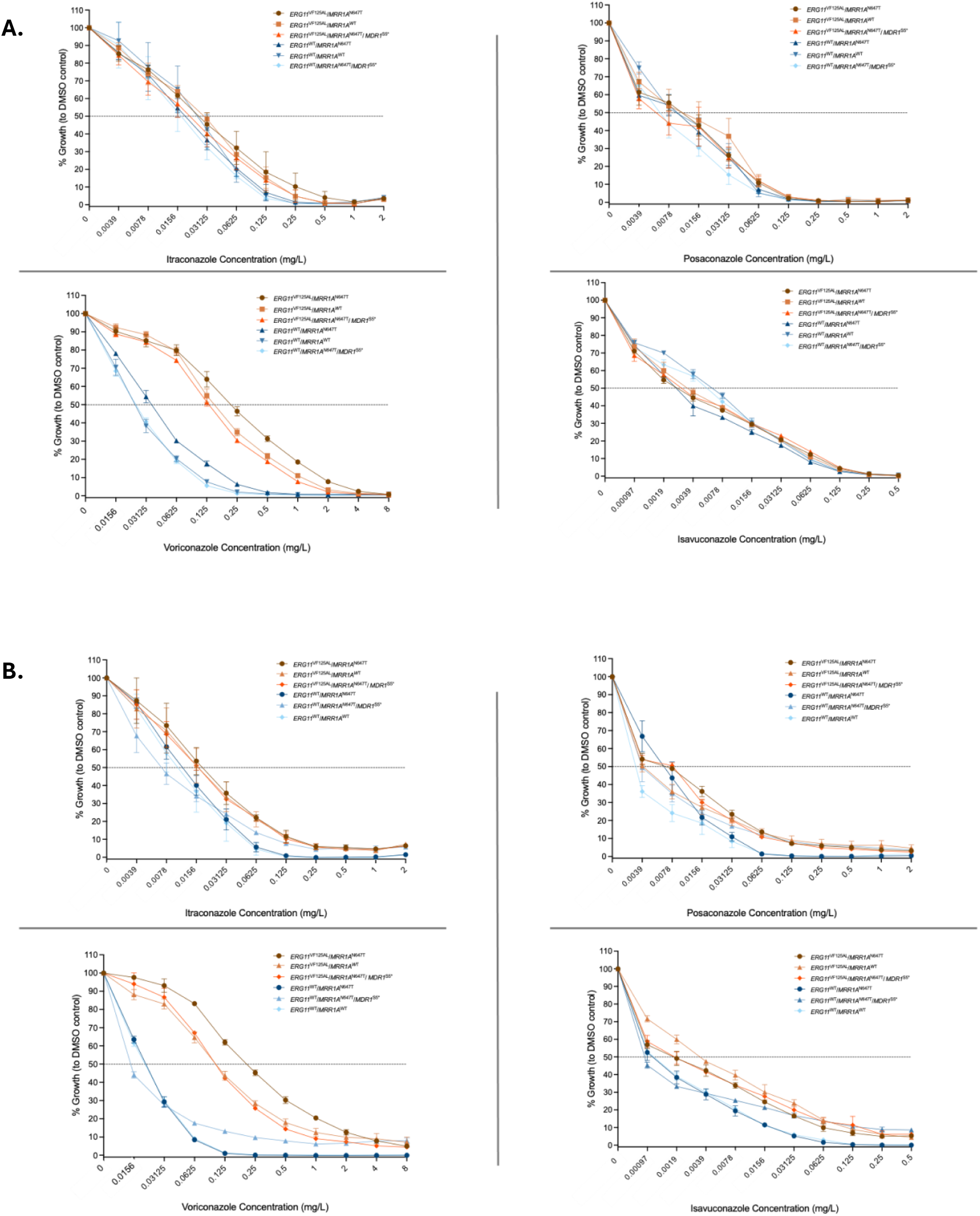
Itraconazole, posaconazole, voriconazole, and isavuconazole MICs of Clade III clinical isolate AR1102 and its derivative strains (A) and Clade III clinical isolate AR0384 and its derivative strains (B) as determined by broth microdilution.

When *ERG11*^VF125AL^ alone was corrected to the wild-type sequence, fluconazole MICs were reduced from 64 µg/mL to 8 µg/mL and 4 µg/mL by broth microdilution and for AR1102 and AR0384, respectively. Correction of both *MRR1A*^N647T^ and *ERG11*^VF125AL^ to their wild-type sequences conferred one additional dilution increase in fluconazole susceptibility to 2 µg/mL for AR1102 and no additional susceptibility for AR0384. Similar changes in susceptibility were observed for voriconazole but not itraconazole, posaconazole, or isavuconazole. These findings indicate that the contribution of *MRR1A*^N647T^ to fluconazole resistance in Clade III isolates is quite modest and that *ERG11*^VF125AL^ is the major driver of fluconazole resistance in this clade. Moreover, the residual fluconazole MIC observed for both isolates is 4 µg/mL when both mutations are edited to their wild-type sequence suggesting the possible involvement of other unknown determinants influencing susceptibility in these isolates.

### The effect of the *MRR1A*^N647T^ mutation on fluconazole susceptibility is dependent upon *MDR1*

In order to determine the contribution of *MDR1* to fluconazole in Clade III isolates, we disrupted this gene in both AR1102 and AR0384. This resulted in a 12– and ten-fold reduction in *MDR1* expression in these isolates, respectively. This also resulted in a one-dilution reduction in fluconazole MIC from 64 µg/mL to 32 µg/mL by broth microdilution but no observable change in MIC by Etest in both isolates phenocopying changes in fluconazole susceptibility observed when *MRR1A* was edited to the wild-type sequence.

We then disrupted *MDR1* in derivatives of AR1102 and AR0384 that retain *MRR1*^N647T^ but that have had *ERG11* edited to the wild-type sequence. This resulted in no additional susceptibility in AR1102 by both broth microdilution and Etest. For AR0384 we observed a one-dilution increase in susceptibility by broth microdilution and shift in MIC from 4 µg/mL to 8µg/mL by Etest. Similar changes were observed for voriconazole by broth microdilution, but not for itraconazole, posaconazole, or isavuconazole. These findings suggest that the modest impact of the *MRR1*^N647T^ mutation on fluconazole and voriconazole susceptibility in Clade III isolates is due to up-regulation of *MDR1*.

## Discussion

The molecular and genetic basis of fluconazole resistance in *Candida* species has proven to be multifactorial and includes mutations in the *ERG11* gene encoding the fluconazole target sterol demethylase [15–17], mutations in the gene encoding the transcriptional regulator of sterol biosynthesis Upc2 [18, 19], mutations in the gene encoding the transcriptional regulator Tac1 which regulates the expression of the genes encoding the ABC transporters Cdr1 and Cdr2 [20, 21], and the transcriptional regulator Mrr1 which regulates the expression of the gene encoding the MFS transporter Mdr1 [9]. Activating mutations in homologs of *MRR1* have likewise been implicated in fluconazole resistance in other *Candida* species such as *C. parapsilosis* and *C. lusitaniae* [22, 23]. Similar mechanisms have been found to be operative in fluconazole resistance in *C. auris*, including mutations in *ERG11* and in the *TAC1* homolog *TAC1B* [6, 7, 15–17].

*C. auris* has three homologs of *MRR1*, designated *MRR1A*, *MRR1B*, and *MRR1C* [10]. A mutation in *MRR1A* leading to the N647T amino acid substitution has been observed in almost all isolates of Clade III [10, 11]. Deletion of *MRR1A* in a Clade III isolate was recently shown to reduce fluconazole and voriconazole MIC as measured by broth microdilution by one dilution from 512 µg/mL to 256 µg/mL and 2 µg/mL to 1 µg/mL, respectively. Deletion of *MRR1A* in a resistant Clade IV isolate carrying the wild-type *MRR1A* allele had no effect on susceptibility [12]. Introduction of the mutation harbored in Clade III isolates into a Clade IV isolate resulted in an increase in fluconazole and voriconazole MIC by two dilutions from 4 µg/mL to 16 µg/mL and 0.06 µg/mL to 0.25 µg/mL, respectively. Deletion of *MDR1* in a susceptible Clade IV isolate had no effect on susceptibility, but its deletion in the presence or absence of the *MRR1A* mutation resulted in the same level of susceptibility [12]. Our observations are consistent with those findings and show that *MRR1*^N647T^ makes a minimal contribution to fluconazole resistance in Clade III isolates. Our work further dissects the effect of *ERG11*^VF125AL^, which is also observed in almost every Clade III isolate described to date and shows that this mutation is responsible for most of the resistance observed in isolates of this clade.

Importantly we found that while these two mutations confer fluconazole resistance and reduced susceptibility to voriconazole in Clade III isolates, they have no effect on the susceptibility of these isolates to isavuconazole or the long chain triazoles itraconazole and posaconazole. This is consistent with the preference of Mdr1 for the short chain triazoles fluconazole and voriconazole in *C. albicans*. This is in contrast to mutations in *TAC1B* leading to increased expression of *CDR1* which confers increased resistance to all of these triazole antifungals. Our findings highlight the possible utility of itraconazole, posaconazole, and isavuconazole in the treatment of infections due to Clade III isolates of *C. auris*.

## Figure Legends

**Supplementary Table 1.**
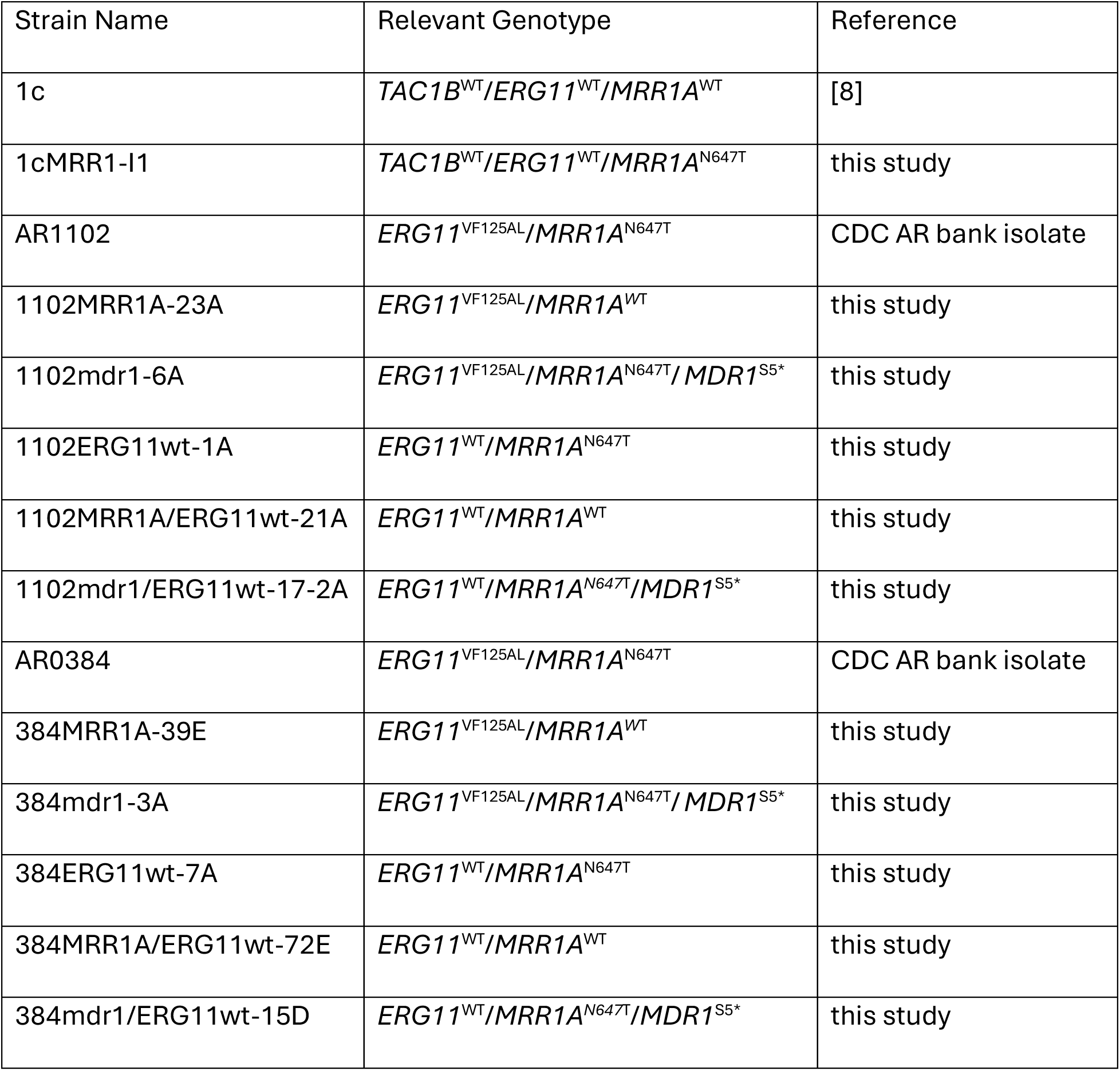
Strains used in this study.

| Strain Name | Relevant Genotype | Reference |
| --- | --- | --- |
| 1c | *TAC1B*<sup>WT</sup>/*ERG11*<sup>WT</sup>/*MRR1A*<sup>WT</sup> | [8] |
| 1cMRR1-I1 | *TAC1B*<sup>WT</sup>/*ERG11*<sup>WT</sup>/*MRR1A*<sup>N647T</sup> | this study |
| AR1102 | *ERG11*<sup>VF125AL</sup>/*MRR1A*<sup>N647T</sup> | CDC AR bank isolate |
| 1102MRR1A-23A | *ERG11*<sup>VF125AL</sup>/*MRR1A*<sup>WT</sup> | this study |
| 1102mdr1-6A | *ERG11*<sup>VF125AL</sup>/*MRR1A*<sup>N647T</sup>/*MDR1*<sup>S5*</sup> | this study |
| 1102ERG11wt-1A | *ERG11*<sup>WT</sup>/*MRR1A*<sup>N647T</sup> | this study |
| 1102MRR1A/ERG11wt-21A | *ERG11*<sup>WT</sup>/*MRR1A*<sup>WT</sup> | this study |
| 1102mdr1/ERG11wt-17-2A | *ERG11*<sup>WT</sup>/*MRR1A*<sup>N647T</sup>/*MDR1*<sup>S5*</sup> | this study |
| AR0384 | *ERG11*<sup>VF125AL</sup>/*MRR1A*<sup>N647T</sup> | CDC AR bank isolate |
| 384MRR1A-39E | *ERG11*<sup>VF125AL</sup>/*MRR1A*<sup>WT</sup> | this study |
| 384mdr1-3A | *ERG11*<sup>VF125AL</sup>/*MRR1A*<sup>N647T</sup>/*MDR1*<sup>S5*</sup> | this study |
| 384ERG11wt-7A | *ERG11*<sup>WT</sup>/*MRR1A*<sup>N647T</sup> | this study |
| 384MRR1A/ERG11wt-72E | *ERG11*<sup>WT</sup>/*MRR1A*<sup>WT</sup> | this study |
| 384mdr1/ERG11wt-15D | *ERG11*<sup>WT</sup>/*MRR1A*<sup>N647T</sup>/*MDR1*<sup>S5*</sup> | this study |

**Supplementary Table 2.**
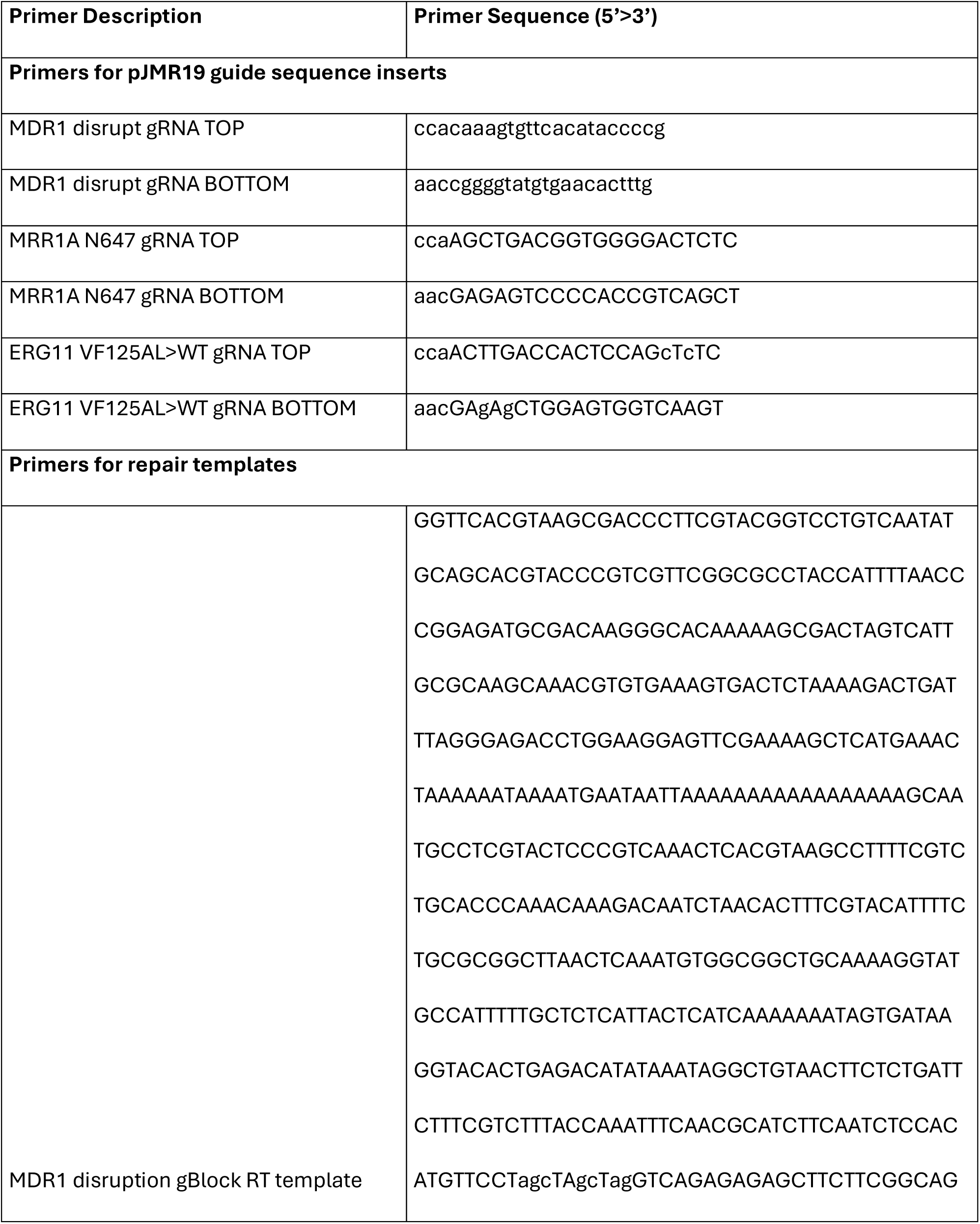
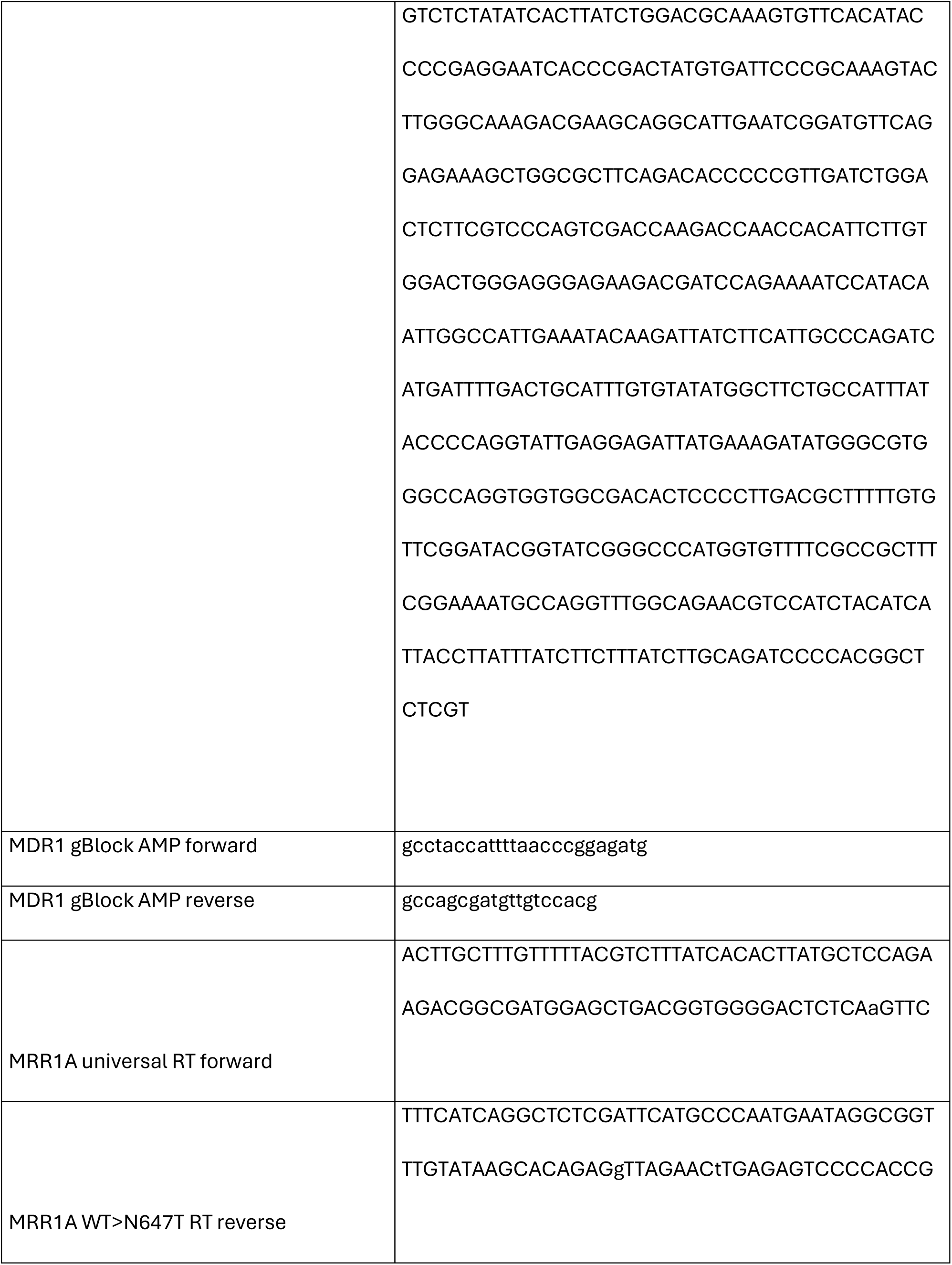
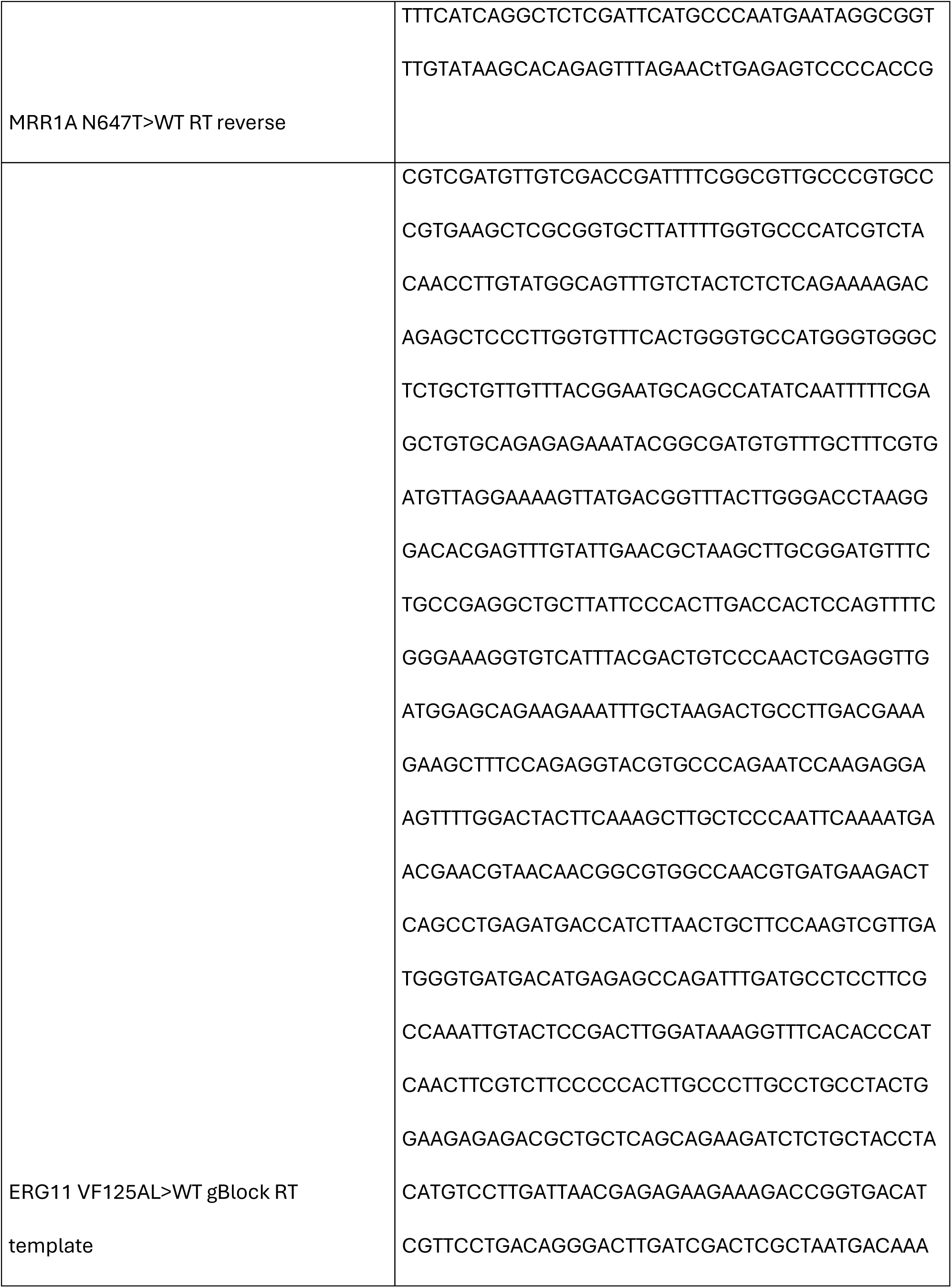
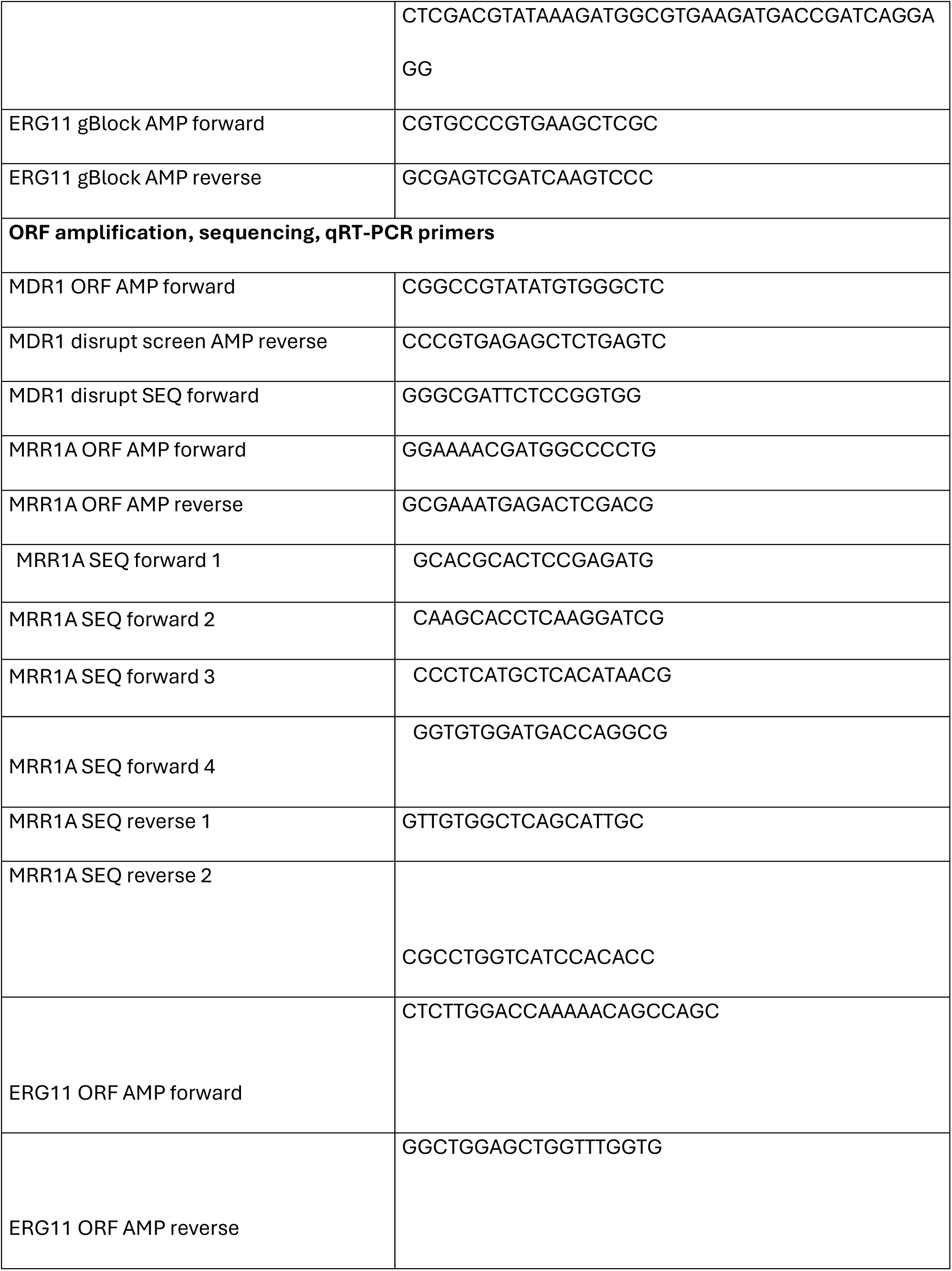
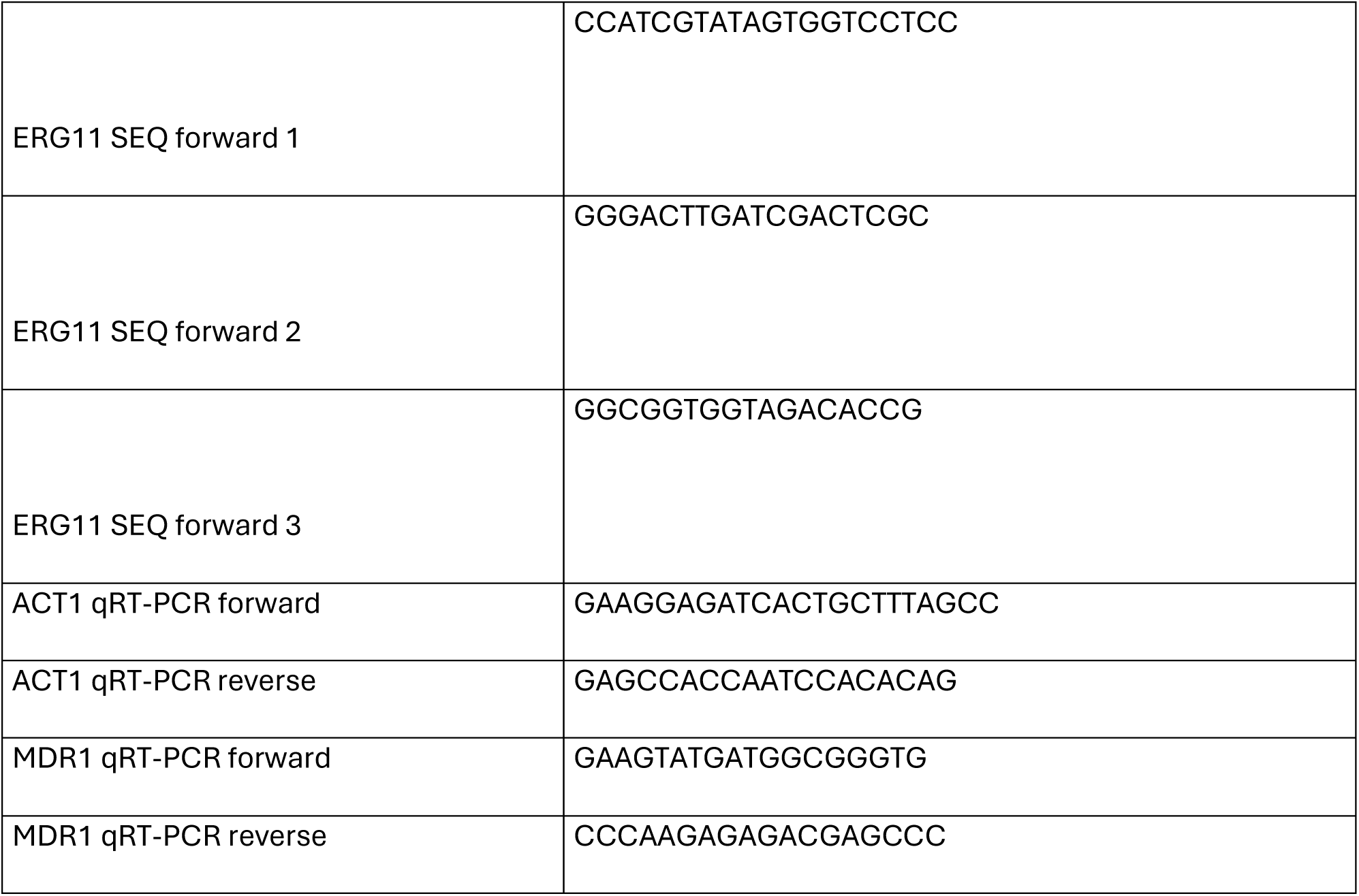
Primers used in this study.

## Notes

### Competing Interest Statement

The authors have declared no competing interest.

